# Parasite escape through trophic specialization in a species flock

**DOI:** 10.1101/098533

**Authors:** Pascal I. Hablützel, Maarten P.M. Vanhove, Pablo Deschepper, Arnout F. Grégoir, Anna K. Roose, Filip A.M. Volckaert, Joost A.M. Raeymaekers

## Abstract

In adaptive radiations species diversify rapidly to occupy an array of ecological niches. In these different niches, species might be exposed to parasites through different routes and at different levels. If this is the case, adaptive radiations should be accompanied by a turnover in parasite communities. How the adaptive radiation of host species might be entangled with such a turnover of parasite communities is poorly documented in nature. In the present study, we examined the intestinal parasite faunas of eleven species belonging to the tribe Tropheini, one of several adaptive radiations of cichlid fishes in Lake Tanganyika. The most parsimonious ancestral foraging strategy among Tropheini is relatively unselective substrate ingestion by browsing of aufwuchs. Certain lineages however evolved more specialized foraging strategies, such as selective combing of microscopic diatoms or picking of macro-invertebrates. We found that representatives of such specialized lineages bear reduced infection with intestinal acanthocephalan helminths. Possibly, the evolution of selective foraging strategies entailed reduced ingestion of intermediate invertebrate hosts of these food-web transmitted parasites. In Tropheini, trophic specialization is therefore intertwined with divergence in parasite infection. We conclude that the study of parasite communities could improve our understanding of host evolution, ecological speciation and the origin of adaptive radiations.

## Introduction

A popular approach to speciation research is to study adaptive radiations during which a lineage evolves rapidly to specialize into an array of distinct ecological niches. Exposure to parasite infection is for a large part determined by the hosts’ niche and ecological divergence can invoke shifts in parasite communities (Knudsen *et al*., 1996; MacColl, 2009a). How parasite communities shift when their host diversifies is an important question in evolutionary ecology that received increased attention in recent years (Eizaguirre *et al*., 2009; MacColl, 2009b; Karvonen & Seehausen, 2012; Vanhove *et al*., 2016). Parasite turnover (e.g. loss of or ‘escape’ from, but also gain of certain parasite taxa) is often considered in a spatial context, where hosts can for example avoid parasites while they invade areas that do not harbour their native parasite communities (‘enemy release hypothesis’ (Keane & Crawley, 2002)). Less documented is ecological or evolutionary escape where changes in host traits lead to parasite escape *in situ* (Chew & Courtney, 1991). In analogy, host species can also become infected with new parasites, e.g. when they adopt a predatory life-style and start feeding on intermediate hosts (Bell & Burt, 1991). Finally, parasite turnover might also be unrelated to diet, but rather follow demographic or phylogenetic divergence of their hosts (Wagner & McCune, 2009; Koblmüller *et al*., 2010; Grégoir *et al*., 2015; Vanhove *et al*., 2015). In the context of adaptive radiations, differentiated parasite communities of incipient host species could impose divergent selection pressures that add up to frequently recognized drivers of speciation such as mate choice or resource competition. Parasites could thereby potentially accelerate and stabilize host divergence (Eizaguirre *et al*., 2009; MacColl, 2009b; Karvonen & Seehausen, 2012). Parasites are ubiquitous in nature, likely comprising more than half of global animal diversity (Windsor, 1998). They are known to exert strong selection on hosts and can be highly host specific (Morand, 2015). Insights in changes of parasite infection upon host diversification are therefore of general relevance and could significantly improve our understanding of adaptive radiations.

The likelihood of parasite infection is, aside from parasite infectivity and host susceptibility, often determined by exposure risk related to habitat use and trophic position. Many parasites have intermediate larval stages infecting prey of their definite (or secondary intermediate or paratenic) hosts (Williams & Jones, 1994). Food-web transmission is therefore a prime infection route, especially for intestinal helminths. At the micro-evolutionary scale, trophic divergence of has been shown to lead to predictable differences in parasite infection among recently diverged species pairs (Karvonen & Seehausen, 2012) or specialized trophic phenotypes (Stutz *et al*., 2014). Similarly, diet has long been recognized as an important predictor of parasite infection at the macro-evolutionary scale (e.g. Drobney *et al*., 1983; Bell & Burt, 1991; Vitone *et al*., 2004; Valtonen *et al*., 2010). Diversification along dietary gradients occurs in many adaptive radiations including the iconic examples of sticklebacks (Schluter, 1996), lake whitefish (Kahilainen *et al*., 2011) and cichlids (Muschick *et al*., 2012). Adaptive radiations should therefore be expected to be accompanied by a predictable turnover in parasite communities. For example, trophic specialization may lead to the avoidance of intermediate hosts as prey items and as such breaking infection routes. The opposite, acquiring new parasite species through new intermediate host prey or other niche-related features (e.g. interactions with other hosts species), is possible too. While it is widely recognized that such turnovers could accelerate and stabilize the process of host species divergence (Eizaguirre *et al*., 2009; MacColl, 2009b), there is a remarkable shortage of empirical research on the interaction between host evolution and parasite infection in adaptive radiations.

The Tropheini tribe comprises one of several adaptive radiations of cichlid fishes from Lake Tanganyika, one of the Great East African Lakes. It currently includes 23 nominal species which occur mostly in sympatry on rocky outcrops in the littoral zone throughout the lake. Phylogenetic relationships among tropheine species are well resolved (Koblmüller *et al*. 2010; Fig. 1A) and the trophic and behavioural ecology of most species has been studied extensively (Kawanabe, Hori & Nagoshi 1997). Tropheini diversified in foraging behaviour and four trophic ecomorphs have been recognized among them: pickers (preying on arthropods), suckers (molluscs), combers (diatoms attached to aufwuchs) and browsers (aufwuchs, mostly consisting of filamentous algae; Yamaoka 1997; Muschick *et al*. 2012; Tada *et al. in press*). Considering the most complete phylogenetic tree of Tropheini (Koblmüller *et al*., 2010), opportunistic browsing of aufwuchs is the most parsimonious ancestral state for the Tropheini radiation from which more specialized foraging strategies have evolved. Browsers are also the most heterogeneous ecomorph regarding their trophic ecology, comprising both specialized aufwuchs-feeders as well as more generalist species supplementing their algae-diet with insects, crustaceans, fish and fish eggs (Muschick *et al*., 2012). Foraging strategies in Tropheini have direct effects on many axes of diversification including morphology and the feeding apparatus (Kawanabe *et al*., 1997; Muschick *et al*., 2012), intestine length (Sturmbauer *et al*., 1992; Wagner *et al*., 2009; Tada *et al., in press*), territorial behaviour (Kawanabe *et al*., 1997) or dispersal capacity (Wagner & McCune, 2009; Koblmüller *et al*., 2010; Grégoir *et al*., 2015; Vanhove *et al*., 2015).

**Fig. 1:**
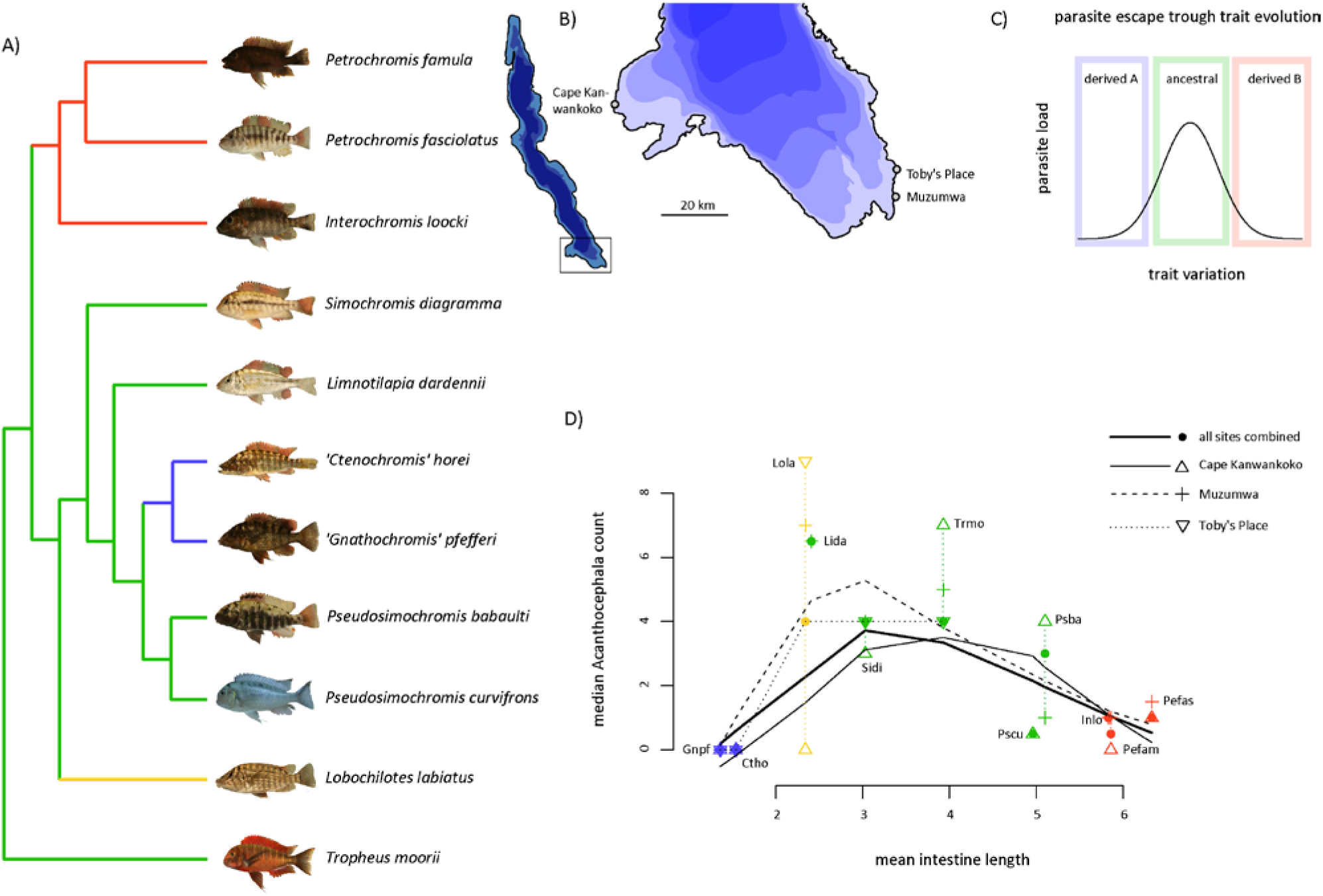
**A)** Cladogram reflecting phylogenetic relationships following Koblmüller *et al*. (2010). Host ecomorph is coded as follows: green = browsers (most parsimonious ancestral state), yellow = suckers, blue = pickers, red = combers. **B)** Sampling sites at the southern shore of Lake Tanganyika in Zambia. **C)** Illustration of theoretical framework of parasite escape through specialization. Hosts with ancestral trait state suffer from high parasite infection. Upon specialization, hosts are less exposed to parasite infection. In the present example, the trait is foraging behaviour with browsing as ancestral and sucking, picking or combing as derived states. **D)** The relative length of intestine (which directly relates on diet and hence foraging ecology) is highly predictive for the abundance of acanthocephalan helminths. Lines are lowess-functions. Species abbreviations include the first two (three) letters of the genus names and species epithets, respectively.

Acanthocephala are the most abundant intestinal parasites of Tropheini (Raeymaekers *et al*., 2013; Hablützel *et al*., 2016). They are typically generalistic parasites with relatively low species diversity and wide host ranges (Vanhove *et al*., 2016). We reported in an earlier study that Acanthocephala infection levels in sympatric *Simochromis diagramma* (Günther 1894) and *Tropheus moorii* Boulenger 1898 co-varied across sites (Hablützel *et al*., 2016), indicating that these hosts (and potentially all Tropheini species) might be infected by the same Acanthocephala species. Acanthocephala exhibit a complex life-cycle with arthropods (commonly amphipods, ostracods or copepods (Williams & Jones, 1994)) serving as intermediate hosts. Infection occurs upon ingestion of the intermediate host (Williams & Jones, 1994), which remains unidentified to date for Lake Tanganyika. We repeatedly observed ostracods in the intestines of several Tropheini species (pers. obs.), making them candidate vectors of Acanthocephala intermediate stages.

During the Tropheini adaptive radiation, evolutionary versatility of the feeding apparatus and novel ecological opportunities allowed species to conquer new positions in the food-web. We therefore hypothesize that trophic specialization within the Tropheini radiation is accompanied by predictable shifts in the intestinal parasite communities. Specifically, we expect that species with little discrimination for the ingested particles (browsers and suckers) more often accidentally swallow intermediate invertebrate hosts than the more specialized pickers and combers. We will test this hypothesis by relating intestine lengths (serving as a univariate proxy for diet) with the abundance of metazoan parasites in eleven species of Tropheini, covering all genera and ecomorphs of this species flock.

## Material & Methods

### Sampling and parasitological screening

Fish were collected in September 2011 and August 2012 at three locations (Cape Kanwankoko (2012): 8° 27′ 8.0″’ S, 30° 27′ 20.0″ E, Muzumwa (2011): 8° 42′ 5.7″ S, 31° 11′ 59.8’’ E and Toby’s Place (2012): 8° 37′ 18.9″ S, 31° 11′ 59.9″ E) at the Zambian shoreline of Lake Tanganyika at a depth of about 0.5 - 3 m (Fig. 1B). Eleven species of Tropheini cichlids encompassing all nine genera were sampled at one, two or three locations respectively (sample sizes in Table 1). At all sites, the collected species occupy the same habitat and are commonly caught in the same net. Between capture and dissection, fish were kept in tanks filled with lake water for at least one night to empty their intestine (which allows for a more reliable parasite count). Keeping fish up to three days in such tanks has little (slight increase for *Gyrodactylus*) or no effect (other parasite taxa) on parasite counts (Raeymaekers et al., 2013). Immediately before dissection, fish were euthanized with an overdose of MS222 and measured to the nearest 0.1 mm (standard length). Intestines were screened for metazoan macro-parasites in the field using a stereomicroscope. The dissection of each fish started with the screening of its integument for monogeneans and crustaceans (copepods, branchiurans, isopods) and any kind of helminthic cyst. The mouth cavity was then inspected for parasitic isopods and branchiurans. Fish were inspected for gill parasites including branchiurans, copepods, bivalves, monogeneans, and any kind of helminthic cyst. To do so, the gills were immediately dissected and stored in 100% ethanol for later processing. Regarding endoparasites, fish were screened for monogeneans, digeneans, acanthocephalans, nematodes, and any kind of helminthic cysts. To do so, stomach, intestines, gall and urinary bladder were dissected immediately after euthanization of the host and inspected in a Petri dish with lake water. Since most host specimen were dissected after they emptied their intestine, the abundance of ostracodes could not be recorded systematically. Processed fish were wrapped in cheese cloth, preserved on formalin and deposited at the Royal Museum for Central Africa (Tervuren, Belgium) as vouchers (samples 2011: collection MRAC B1.23; samples 2012: collection MRAC B2.38).

**Table 1:** Sampling sizes and median Acanthocephala abundance for eleven Tropheini species. Dashes at Cape Kanwankoko and Muzumwa indicate that the species was very rare (and not necessarily absent) at the respective sampling site. Dashes at Toby's place reflect limited sampling efforts due to different initial study aims. Full names of species can be found in Fig. 1A.

| Species | Sample size |  |  |  | Median Acanthocephala abundance |  |  |  |
| --- | --- | --- | --- | --- | --- | --- | --- | --- |
|  | Cape Kanwankoko | Muzumwa | Toby's Place | all sites | Cape Kanwankoko | Muzumwa | Toby's Place | all sites |
| Ctho | 5 | 6 | 10 | 21 | 0 | 0 | 0 | 0 |
| Gnpf | 5 | 5 | 10 | 20 | 0 | 0 | 0 | 0 |
| Lida | 0 | 6 | 0 | 6 | – | 6.5 | – | 6.5 |
| Lola | 5 | 6 | 10 | 21 | 0 | 7 | 9 | 4 |
| Inlo | 0 | 6 | 0 | 6 | – | 1 | – | 1 |
| Pefam | 5 | 7 | 0 | 12 | 0 | 1 | – | 0.5 |
| Pefas | 5 | 6 | 0 | 11 | 1 | 1.5 | – | 1 |
| Psba | 5 | 5 | 0 | 10 | 4 | 1 | – | 3 |
| Pscu | 6 | 0 | 0 | 6 | 0.5 | – | – | 0.5 |
| Sidi | 5 | 31 | 35 | 71 | 3 | 4 | 4 | 4 |
| Trmo | 5 | 89 | 86 | 180 | 7 | 5 | 4 | 4 |
| Total | 46 | 167 | 151 | 364 |  |  |  |  |

### Intestine length as an univariate proxy for diet

Specialization in foraging ecology among Tropheini species has repeatedly and strongly been associated with evolutionary changes in intestine length (Sturmbauer *et al*., 1992; Wagner *et al*., 2009; Tada *et al., in press*). Pickers feed on readily digestible prey and, in line with the costly tissue hypothesis (Tsuboi *et al*., 2016), evolved short intestines. In contrast, combers and some specialized browsers (both species of *Pseudosimochromis* Nelissen 1977 represented in our sampling) have long intestines that can digest low quality diet such as diatoms or filamentous algae. Mean intestine length (as a proportion of host standard length) per species was obtained from Tada *et al*. (*in press*).

### Statistical analyses

We used a generalized linear model (GLM) to describe the effects of host species, sampling site, host size (standard length) and host sex (male, female or immature) on abundance (count of parasite specimens per host individual) of acanthocephalan helminths, and the gill-infecting ectoparasite genera *Cichlidogyrus* Paperna 1960 (Monogenea, Ancyrocephalidae) and *Ergasilus* von Nordmann 1832 (Copepoda, Ergasilidae). Other parasites (Nematoda, Digenea and unidentified helminthic cysts) were found in the intestines in low numbers (overall prevalence < 0.05), preventing the application of statistical models. The effect of sampling year was confounded with sampling site and was not included in the model. We have shown earlier that parasite infection was relatively stable between the two sampling years in one host species (*T. moorii;* Raeymaekers *et al*., 2013). Since we were interested in how far the species effect varies among sites, we ran the model a second time after adding a species x site interaction effect. Abundance was fit on a GLM assuming a Poisson distribution of parasite counts. Analysis of variance was conducted using type II sums of squares.

The association between species-level variation of intestine length and median abundance of acanthocephalans, *Cichlidogyrus* or *Ergasilus* per host species and per site was analysed in a second-order polynomial regression model. Sampling site was included as a random effect. Since the distribution of parasite counts was not normal (few host individuals had many parasites) and the relationship between Acanthocephala counts and intestine was right-tailed (Fig. 1D), both response and predictor variables were log-transformed prior to statistical analyses. We repeated the analysis with a reduced dataset from which all host species with less than 10 specimens (*Interochromis lookii* (Poll 1949), *Limnotilapia dardennii* (Boulenger 1899) and *Pseudosimochromis curvifrons* (Poll 1942)) were excluded in order to assess whether our analysis was sensitive for the limited sample size of some of the host species. To test the hypothesis that parasites replace each other across host taxa, we conducted statistical tests for Pearsons’s product-moment correlations among median abundances of the three most abundant parasite groups. All analyses were conducted in R v.3.3.0 (R Development Core Team, 2011).

## Results

Parasites infecting every species included intestinal acanthocephalans, the ancyrocephalid monogenean *Cichlidogyrus* and the copepod *Ergasilus* on the gills. Parasites which were not present on every single host species included the gyrodactylid monogenean *Gyrodactylus* on skin and fins, intestinal nematodes, the monogenean *Urogyrus* in the urinary bladder, branchiurans in the gill cavity or on the opercula, intestinal digeneans, and a number of unidentified helminthic cysts in skin, fin or gill tissue. Acanthocephalans (found in all host species; median abundance: 0-6.5; Fig. 1D) dominated the intestinal parasite fauna while nematodes (7 host species; median abundance: 0), digeneans (2 host species; median abundance: 0) and helminthic cysts (3 host species; median abundance: 0) were observed sporadically and in low numbers (Appendix S1 Table S1). Host species was the strongest predictor of abundance of all parasite groups, while sampling sites, host size, host sex and the interaction between species and site had minor, although significant (except for host size on Acanthocephala and host sex on *Ergasilus*), effects (Table 2).

**Table 2:** Results of the generalized linear model for host species-level variation in abundance of Acanthocephala, *Cichlidogyrus* and *Ergasilus* accounting for confounding effects of sampling site, host standard length and host sex.

| Acanthocephala |  |  |  |  |
| --- | --- | --- | --- | --- |
| Effect | LR $X^2$ | Num DF | Den DF | Pr(> $X^2$) |
| Species | 308.89 | 10 | 149 | < 0.001 |
| Site | 37.44 | 2 | 149 | < 0.001 |
| SL | 1.54 | 1 | 149 | 0.214 |
| Sex | 16.89 | 2 | 149 | < 0.001 |

| Effect | LR $X^2$ | Num DF | Den DF | Pr(> $X^2$) |
| --- | --- | --- | --- | --- |
| Species | 308.89 | 10 | 149 | < 0.001 |
| Site | 37.44 | 2 | 149 | < 0.001 |
| SL | 0.57 | 1 | 149 | 0.449 |
| Sex | 29.93 | 2 | 149 | < 0.001 |
| Species x Site | 81.76 | 11 | 149 | < 0.001 |

***Cichlidogyrus***
| Effect | LR $X^2$ | Num DF | Den DF | Pr(> $X^2$) |
| --- | --- | --- | --- | --- |
| Species | 5841.70 | 10 | 134 | < 0.001 |
| Site | 394.91 | 2 | 134 | < 0.001 |
| SL | 944.42 | 1 | 134 | < 0.001 |
| Sex | 67.00 | 2 | 134 | < 0.001 |
| Species | 5841.70 | 10 | 134 | < 0.001 |
| Site | 394.91 | 2 | 134 | < 0.001 |
| SL | 592.77 | 1 | 134 | < 0.001 |
| Sex | 30.55 | 2 | 134 | < 0.001 |
| Species x Site | 356.72 | 11 | 134 | < 0.001 |

***Ergasilus***
| Effect | LR $X^2$ | Num DF | Den DF | Pr(> $X^2$) |
| --- | --- | --- | --- | --- |
| Species | 612.24 | 10 | 133 | < 0.001 |
| Site | 108.71 | 2 | 133 | < 0.001 |
| SL | 164.02 | 1 | 133 | < 0.001 |
| Sex | 2.20 | 2 | 133 | 0.333 |
| Species | 612.24 | 10 | 133 | < 0.001 |
| Site | 108.71 | 2 | 133 | < 0.001 |
| SL | 110.1 | 1 | 133 | < 0.001 |
| Sex | 1.22 | 2 | 133 | 0.543 |
| Species x Site | 90.91 | 11 | 133 | < 0.001 |

Intestine length significantly predicted median Acanthocephala abundance (Table 3). The association was curvilinear with species with short or long intestines bearing the lowest number of Acanthocephala (Table 3; Fig. 1D). The polynomial regression term remained significant after removing three host species with low sample sizes (p = 0.004). Pickers with short intestines had zero median abundance, although Acanthocephala could occasionally be observed in all host species (Appendix S1 Fig. S1). Suckers and three genera (*Limnotilapia* Regan 1920, *Simochromis* Boulenger 1898 and *Tropheus* Boulenger 1898) of browsers with intermediate intestine length showed the highest Acanthocephala infection (median abundance: 4-6.5). The two browser species of the genus *Pseudosimochromis* with rather long intestines were infected with relatively low numbers of Acanthocephalans (median abundance: 0.5-3). Finally, combers also showed low Acanthocephala abundance (median abundance: 0.5-1). Intestine length was not significantly correlated with *Cichlidogyrus* and *Ergasilus* counts (Table 3). The relationships did not change upon removal of three host species with low sample sizes (p = 0.579 and p = 0.184). We further found that parasite groups did not replace each other across host taxa (Acanthocephala vs. *Cichlidogyrus:* correlation coefficient = −0.143, p-value = 0.676; Acanthocephala vs. *Ergasilus:* correlation coefficient = −0.340, p-value = 0.307; *Cichlidogyrus* vs. *Ergasilus:* correlation coefficient = 0.407, p-value = 0.214).

**Table 3:** Results of the general linear mixed-effect model for the relationship between intestine length (predictor) and median parasite count (response). Sampling site was included as a random effect. Wald X^2^-tests were used to assess the statistical significance of the linear model fit or of the improvement of the application of a second-order polynomial function, respectively.

| Parasite | Model | Df | AIC | BIC | logLik | Deviance | Test | $\chi^2$ | X Df | Pr(>$\chi^2$) |
| --- | --- | --- | --- | --- | --- | --- | --- | --- | --- | --- |
| Acanthocephala | linear | 6 | 128.0 | 135.0 | -58.0 | 116.0 | linear | 0.25 | 1 | 0.615 |
|  | polynomial | 10 | 117.6 | 129.4 | -48.8 | 97.6 | linear vs. polynomial | 18.3 | 4 | 0.001 |
| *Cichlidogyrus* | linear | 6 | 225.8 | 232.8 | -106.9 | 213.8 | linear | 0.74 | 1 | 0.390 |
|  | polynomial | 10 | 231.0 | 242.8 | -105.5 | 211.0 | linear vs. polynomial | 2.7 | 4 | 0.602 |
| *Ergasilus* | linear | 6 | 152.8 | 159.8 | -70.4 | 140.8 | linear | 0.53 | 1 | 0.467 |
|  | polynomial | 10 | 154.0 | 165.8 | -67.0 | 134.0 | linear vs. polynomial | 6.7 | 4 | 0.150 |

## Discussion

We hypothesized that diversification in foraging ecology could be accompanied by shifts in (intestinal) parasite communities as found in several sympatric species pairs (Knudsen *et al*., 1997; MacColl, 2009a). Using the adaptive radiation of the Lake Tanganyika cichlid tribe Tropheini as a model, we found that the abundance of trophically transmitted acanthocephalan helminths was predicted by inter-specific variation in intestine length, which itself is strongly correlated with differentiation in foraging strategy and diet (Sturmbauer *et al*., 1992; Wagner *et al*., 2009; Tada *et al., in press*). This observation was not paralleled by ectoparasites. We discuss to which extent trophic diversification and parasite infection are intertwined and how this interplay might affect the hosts’ adaptive radiation.

### Foraging ecology predicts parasite infection

Tropheini species could escape their acanthocephalan parasites twice by evolving specialized feeding strategies (although escape is incomplete, since both pickers and combers may be infested with low numbers of Acanthocephala; Figs 1C and 1D). Browsers (the most parsimonious ancestral ecomorph) shear filamentous algae *in toto* from the substrate (Yamaoka, 1997), along with the associated micro-invertebrate fauna (thus including the putative intermediate host of acanthocephalans). The grazing species *Petrochromis* spp. and *Interochromis loocki*, in contrast, are specialized diatom feeders who comb their food from filamentous algae (Yamaoka, 1997). They are therefore able to selectively ingest tiny particles (thus excluding the putative intermediate host, which measures around 1 mm, while diatoms range from about 0.002-0.2 mm). The picker-lineage encompassing *'Ctenochromis' horei* (Günther 1894) and ‘*Gnathochromis*’ *pfefferi* (Boulenger 1898) evolved into selective predators of insect larvae and larger crustaceans (e.g. shrimps (Muschick *et al*., 2012)) that probably do not carry Acanthocephala larval stages. The sucker *Lobochilotes labiatus* (Boulenger 1898) also preys on macro-invertebrates (mainly molluscs (Colombo *et al*., 2013), which are not known as hosts of Acanthocephala (Williams & Jones, 1994)). However, due to its sucking feeding behaviour, the species is (similar to browser species) relatively indiscriminate about the ingested items (Muschick *et al*., 2012). Indeed, we found, on average, high infection with acanthocephalan parasites in *L. labiatus*.

The relationship between parasite infection and host trophic ecology might be confounded by geographic variation in parasite abundance and host-parasite co-evolutionary interactions (Bell & Burt, 1991; Stutz *et al*., 2014). Acanthocephala infections do indeed vary across the study area (Raeymaekers *et al*., 2013; Hablützel *et al*., 2016; this study) but we found the confounding effect of geography to be of little importance compared to the main host species effect. Ultimately, parasite load will not only be influenced by ecological (exposure to propagules) but also evolutionary (parasite virulence and host susceptibility) factors. Acanthocephalans are known to interact with the immune system, although pathological effects are typically only observed upon massive infection (Paperna, 1996). Laboratory experiments provided empirical evidence for heritable variation in susceptibility to Acanthocephala infection in sticklebacks (Mazzi & Bakker, 2003), indicating that different degrees of resistance could explain variation in infection intensities among host species. Resistance to parasite infection might come at an immunological cost (Råberg *et al*., 2009) that trades off against the parasite burden, favouring tolerance towards the parasite if its virulence is low. In at least one species of Tropheini cichlids (*T. moorii*), Acanthocephala infection has little or no effect on host body condition, indicating some degree of tolerance evolution (Hablützel *et al*., 2014). The selective pressure for tolerance or resistance evolution might be expected to be itself related to exposure risk. Species suffering from high exposure should therefore experience the strongest selection pressure to become resistant (or tolerant), a hypothesis that is not unlikely, but cannot be tested with the current data.

### Parasite infection: an understudied dimension of adaptive radiations

Specialization in foraging ecology is one of the most prominent processes in adaptive radiations. Species divergence in this context is often considered a consequence of character displacement due to resource competition (Schluter, 1994). The observation that parasite infection is inherently intertwined with trophic diversification adds an understudied dimension to this process. Speciation models and field studies suggest that trophic niche partitioning might be plastic at first and becomes heritable upon genetic divergence of the incipient species (Pfennig *et al*., 2010). Evolutionary escape from parasites might accelerate and stabilize this process in two ways. First, the cost of adaptation to new food sources might be compensated by parasite escape. Second, immunity gene pools might diverge among incipient host species (Eizaguirre *et al*., 2012) under both resistance or tolerance scenarios. Dietary versatility through phenotypic plasticity might become costly upon immunogenetic divergence, since neither of the diverging host lineages will be immunogenetically adapted to the parasite community that is associated with the alternative foraging strategy. Certainly, the strong co-variance between parasite community variation and niche divergence of their hosts highlights an understudied component of adaptive radiations.

## Acknowledgements

We thank J. Bamps and S. Camey for help with fieldwork and parasite dissections. Furthermore, we thank L. Makasa, D. Sinyinza, G. Sheltons, C. Sturmbauer, W. Salzburger, W. Mubita and the staff of the Lake Tanganyika Research Station in Mpulungu (Zambia) for help with fieldwork and logistics. Research was supported by the Research Foundation – Flanders (FWO grant project G.0553.10), the Flemish Interuniversity Council (VLIR) and the KU Leuven Research Fund project PF/2010/07. PIH was partially supported by the Janggen-Pöhn-Stiftung (St. Gallen, Switzerland). MPMV is partly supported by the Czech Science Foundation, Project no. P505/12/G112 (European Centre of Ichthyoparasitology (ECIP) – Centre of excellence). AFG is a PhD fellow of the Research Foundation – Flanders. JAMR received a EU Marie Sktodowska-Curie Fellowship (IEF 300256).

